# Receptor Binding Specificity of a Bovine A(H5N1) Influenza Virus

**DOI:** 10.1101/2024.07.30.605893

**Authors:** Pradeep Chopra, Caroline K. Page, Justin D. Shepard, Sean D. Ray, Ahmed Kandeil, Trushar Jeevan, Andrew S. Bowman, Ali H. Ellebedy, Richard J. Webby, Robert P. de Vries, S. Mark Tompkins, Geert-Jan Boons

## Abstract

Outbreaks in the US of highly pathogenic avian influenza virus (H5N1) in dairy cows have been occurring for months creating new possibilities for direct contact between the virus and humans. Eisfeld *et al*. examined the pathogenicity and transmissibility of a bovine HPAI H5N1 virus isolated from New Mexico in a series of *in vitro* and *in vivo* assays. They found the virus has a dual human- and avian virus-like receptor-binding specificity as measured in a solid phase glycan binding assay. Here, we examined the receptor specificity of a bovine HPAI H5N1 virus (A/bovine/OH/B24OSU-432/2024, H5N1, clade 2.3.4.4b) employing four different assays including glycan array technology, bio-layer interferometry (BLI), a solid phase capture assay and hemagglutination of glycan remodeled erythrocytes. As controls, well characterized avian (A/Vietnam/1203/2004, H5N1, clade 1) and human (A/CA/04/2009, H1N1) IAVs were included that bind α2,3- and α2,6-sialosides, respectively. We found that A/bovine/OH/B24OSU-432/2024 preferentially binds to “avian type” receptors (α2,3-sialosides). Furthermore, sequence alignments showed that A/bovine has maintained amino acids in its HA associated with α2,3-sialoside (avian) receptor specificity. We conclude that while we find no evidence that A/bovine has acquired human virus receptor binding specificity, ongoing efforts must be placed on monitoring for this trait.

---

Highly pathogenic avian influenza (HPAI) viruses of the H5Nx A/goose/Guangdong/1/96 lineage, which emerged in China in 1996, are a cause of pandemic concern because of their frequent spill over into mammals^1^. In 2021, new H5N1 viruses belonging to the 2.3.4.4b hemagglutinin (HA) phylogenetic clade appeared and have become endemic in wild birds in multiple continents, causing frequent infections in poultry. These avian viruses have also been detected in several mammalian species including humans, elevating concerns regarding the pandemic potential of these viruses^2^.

In early 2024, infections with HPAI H5N1 clade 2.3.4.4b viruses were detected in dairy cattle in Texas (USA) and these viruses have now spread and been confirmed in farms in thirteen states in the US^3,4^. Clinical signs are mainly observed in lactating cows and include decreased feed intake, altered faecal consistency, respiratory distress, and decreased and abnormal milk production. Viral staining of tissues has revealed a distinct tropism for epithelial cells lining the alveoli of the mammary gland. Furthermore, milk collected from affected cows is commonly contaminated with high loads of infectious virus. There is evidence for efficient cow-to-cow transmission, and genetic analyses of viruses from dairy cows, birds, domestic cats, and a raccoon from affected farms indicate interspecies transmissions^5^. Four dairy cow farmworkers have become infected by bovine H5N1 viruses displaying clinical symptoms primarily of conjunctivitis with some respiratory signs reported^6^.

Eisfeld *et al*. recently examined the pathogenicity and transmissibility of a bovine HPAI H5N1 virus isolated from New Mexico in a series of *in vitro* and *in vivo* assays^7^. Of note, these studies found the virus had a dual human- and avian virus-like receptor-binding specificity as measured in a solid phase glycan binding assay. The receptor binding preference of HA of influenza A viruses (IAV) is a major determinant of host range^8,9^. Avian viruses preferentially bind α2,3-linked sialic acids, which are found in the duck enteric and chicken upper respiratory tract, whereas human IAVs recognize α2,6-linked sialic acids, which are found in the upper respiratory tract of humans. In the lower respiratory tract and conjunctiva of humans, “*avian-type*” receptors are also expressed allowing for infection by avian viruses but limiting onward transmission. The human viruses of the H1, H2, and H3 subtypes that caused pandemics in 1918, 1957, and 1968, respectively, were of avian origin, and their HA adapted to recognizing α2,6-linked sialic acids. It is anticipated that gaining ‘*human-type*’ receptor specificity is required for sustained human-to-human transmission of avian influenza viruses, including the bovine H5N1 virus.

Here, we examined the receptor specificity of a bovine HPAI H5N1 virus (A/bovine/OH/B24OSU-432/2024, H5N1, clade 2.3.4.4b) employing four different assays including glycan array technology, bio-layer interferometry (BLI), a solid phase capture and hemagglutination of glycan remodeled erythrocytes. The amino acid sequence of the HA of A/bovine/OH/B24OSU-432/2024 is identical to that of the virus examined by Eisfeld *et al*. (Table S1). As controls, we included well characterized avian (A/Vietnam/1203/2004, H5N1, clade 1) and human (A/CA/04/2009, H1N1) IAVs that bind α2,3- and α2,6-sialosides, respectively. We found that A/bovine/OH/B24OSU-432/2024 preferentially binds to “avian type” receptors (α2,3-sialosides).

Previously, we synthesized a panel of glycans that resemble structures found on respiratory tissues^10^. The panel includes *N*-glycans that have multiple consecutive *N*-acetyl-lactosamine (Gal(β1,4)GlcNAc, LacNAc) repeating units that can either be unmodified (**A**-**C**) or capped by α2,3-linked (**D**-**N**) or α2,6-linked sialosides (**O**-**X**). Also included are several core-2 *O*-linked glycopeptides that are devoid of sialic acid (**1-2**) or have an α2,3-sialyl LacNAc moiety (**3**) that can be further modified by fucosylation (**4**) and sulfation (**5**) (Fig. 1A). Such glycans have been observed in mucins and their fucosylation to give sialyl Lewis^x^ is a modification tolerated by HPAI of the clade 2.3.4.4 due to K222Q and S227R substitutions^11,12^. Furthermore, sulfation of GlcNAc can affect the binding properties of IAVs and, for example, some H5/H7 subtype IAVs prefer Neu5Acα2,3Galβ1,4(6-SO_3_)GlcNAc^13^. The synthetic *N*-glycans and glycopeptides were printed on amine-reactive NHS activated glass slides and quality control was performed using biotin-labeled plant lectins such as ECA, MAL-I, and SNA (Fig. S1). Live or β-propiolactone (BPL)-inactivated whole viruses were applied to the microarray in the presence of oseltamivir (10 μM), and detection of binding was determined using a pan-HA anti-stalk antibody (1A06). The attraction of using whole viruses for receptor binding studies is that multiple copies of HA are presented, thereby providing proper multivalency, which is important for receptor specificity. Microarray screening of A/bovine/OH/B24OSU-432/2024 (BPL-inactivated) showed binding only to α2,3-sialosides (Fig. 1B). Fine specificities were observed for A/bovine/OH/B24OSU-432/2024 preferring bi-antennary *N*-glycans having extended LacNAc moieties at both antennae modified as α2,3-sialosides (**G**-**K, M** and **N**). Compounds with a sialoside at only one antenna (*e*.*g*. **D, E, F**) gave lower responsiveness. A/bovine/OH/B24OSU-432/2024 also recognizes *O*-glycans that have sialyl LacNAc moieties (**3**) that can be further modified by an α1,3-fucoside (**4**, SLe^x^) or 6-*O*-sulfate (**5**). The evolutionarily earlier A/Vietnam/1203/2004 (H5N1) virus (BPL-inactivated) had a more restricted binding pattern and did not tolerate fucosylation, such as in compound **5**. These observations are consistent with the HA K222Q and S227R substitutions in the bovine virus, creating a binding pocket for fucose^11^. As expected, A/CA/04/2009 (H1N1) bound only to *N*-glycans having a 2,6-sialoside and displayed a preference for compounds that have at least an extended 2,6-sialoside at the α(2,3)-antennae.

**Fig. 1.**
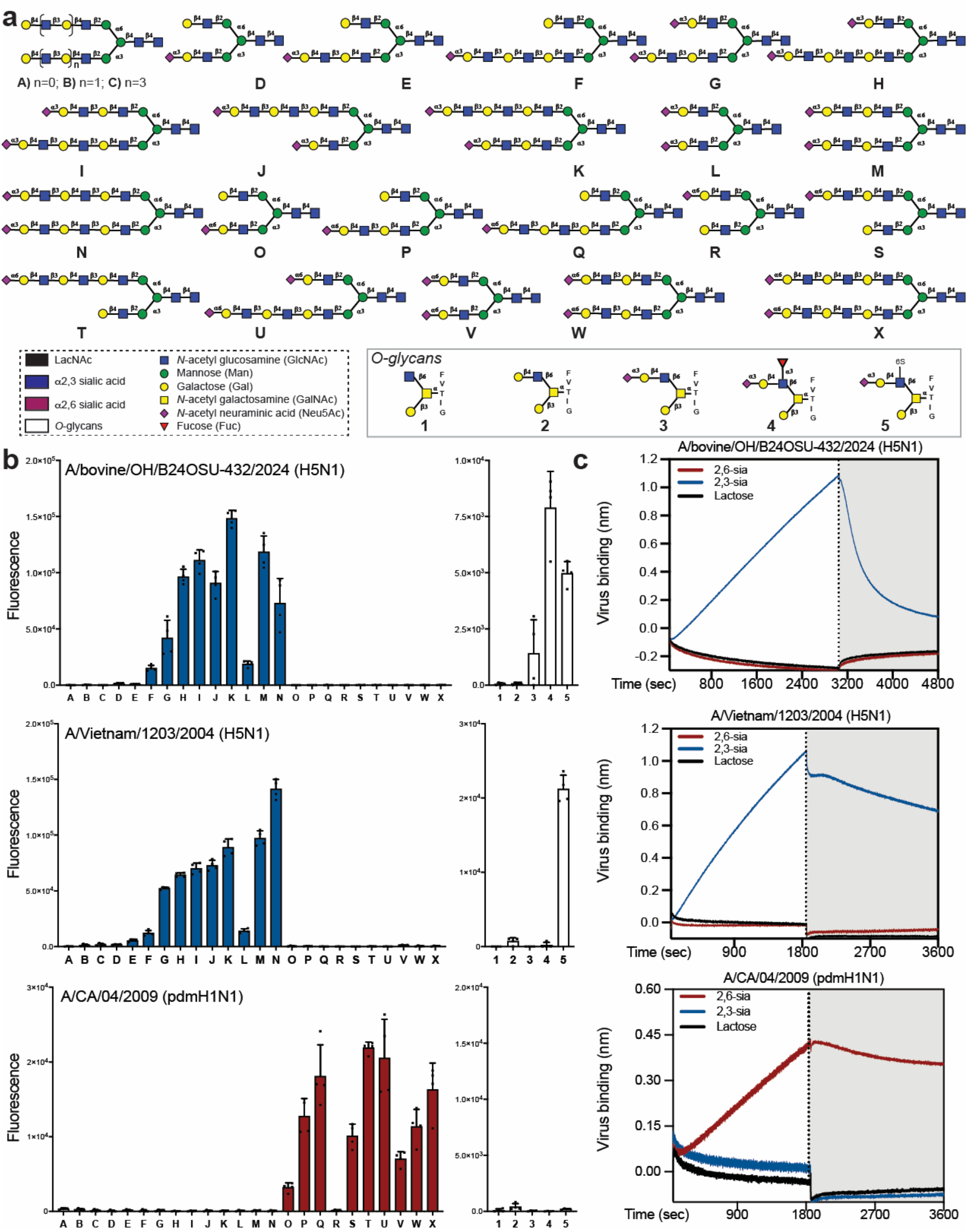
Receptor binding specificity of bovine H5N1. (a) Structures of glycans printed on the microarray. All *N*-glycans (**A**-**X**) have an α-amine at the reducing end asparagine moiety and *O*-glycans (**1**-**5**) have a *N*-terminal α-amine. (b) Glycan array binding analysis of the receptor binding specificities of H5N1 and H1N1 viruses. The fluorescence signals for each glycan are presented as mean ± SD (n = 4). (c) Bio-layer interferometry (BLI) sensorgrams demonstrating the sialic acid linkage-specificity of H5N1 and H1N1 viruses. Virus binding (association, in the presence of Oseltamivir,) and virus elution (dissociation, in the absence of Oseltamivir, gray highlighted) are shown. Representative data of at least two replicate experiments are shown.

Next, the interaction of the viruses with glycans was examined by bio-layer interferometry (BLI), which is a label-free technology for quantifying biomolecular interactions in real time^14^. Streptavidin coated BLI sensors were loaded with biotinylated poly(acrylamide) (PAA) polymers that were modified by α2,3-sialyl-LacNAc, α2,6-sialyl-LacNAc or lactose. Next, the sensors were dipped into a buffer containing A/bovine/OH/B24OSU-432/2024 (H5N1), A/Vietnam/1203/2004 (H5N1) and A/CA/04/2009 (H1N1) in the presence of a neuraminidase inhibitor (Oseltamivir, 10 μM) (Fig. 1C). The H5 viruses bound to probes modified by α2,3-sialosides, whereas no binding was observed for probes derivatized by α2,6-sialosides even after an extended incubation time. A/CA/04/2009 (H1N1) bound only to sensors modified by α2,6-sialosides. None of the viruses bound to negative control sensors functionalized with lactose.

We also performed a solid phase binding assay in which the viruses were captured on fetuin-coated microtiter plates and then probed for binding using biotinylated PAA polymers modified by α2,3-sialyl-LacNAc, α2,6-sialyl-LacNAc or lactose in the presence of Oseltamivir, followed by incubation with horseradish peroxidase (HRP) conjugated streptavidin. A/bovine/OH/B24OSU-432/2024 (H5N1) and A/Vietnam/1203/2004 (H5N1) bound only to α2,3-sialosides and A/CA/04/2009 (H1N1) to α2,6-sialosides (Fig. S2).

Hemagglutination, in which HA of viral particles bind to sialic acid receptors of red blood cells (RBCs), is commonly employed to detect IAVs. RBCs express a plethora of glycans, and therefore, viruses with different receptor specificities can hemagglutinate RBCs. We have developed an exo-enzymatic cell surface glycan remodelling strategy to install specific glycan receptors on fowl erythrocytes^15^. In this study, we treated fresh turkey erythrocytes with *Arthrobacter ureafaciens* sialidase to remove all sialic acid receptors followed by resialylation with the α2,3-sialyltransferase from *Pasteurella multocida* sialyltransferase (Pmst1 M144D) or ST6 β-galactoside α-2,6-sialyltransferase 1 (ST6Gal1) in the presence of CMP-Neu5Ac to install α2,3- or α2,6-linked sialosides, respectively. The untreated RBCs, those lacking sialic acid (sialidase treated), or those displaying α2,3- or α2,6-linked sialosides were examined for hemagglutination by the various IAVs. A/bovine/OH/B24OSU-432/2024 and A/Vietnam/1203/2004 (H5N1) agglutinated untreated and α2,3-sialoside modified erythrocytes (Table 1). On the other hand, A/CA/04/2009 (H1N1) agglutinated the untreated and α2,6-sialoside modified erythrocytes. Experiments using live A/bovine/OH/B24OSU-432/2024 (H5N1) in an enhanced BSL3 facility gave similar results although some low-level binding to α2,6-sialoside modified RBC was detected.

**Table 1.** Hemagglutination (HA) titers of influenza A viruses with remodelled erythrocytes. Glycan remodelled turkey erythrocytes were used for HA assays with either live or BPL-inactivated A/California/04/2009, A/Vietnam/1203/2004, and A/Bovine/Ohio/B24OSU-432/2024 in BSL2 (experiment 1 and 2) and enhanced BSL3 (experiment 3) laboratories. Freshly remodelled erythrocytes were prepared for each independent experiment. *Data is reported as means of two technical replicates.

| Independent Experiment | Virus | Erythrocyte Treatment |  |  |  |
| --- | --- | --- | --- | --- | --- |
| | | Untreated | Sialidase | α2,3-Sia | α2,6-Sia |
| 1 | A/CA/04/2009 (Live) | 32 | 0 | 0 | 32 |
|  | A/Vietnam/1203/2004 (BPL) | 32 | 0 | 32 | 0 |
|  | A/bovine/OH/B24OSU-432/2024 (BPL) | 32 | 0 | 32 | 0 |
| 2 | A/CA/04/2009 (Live) | 32 | 0 | 0 | 32 |
|  | A/Vietnam/1203/2004 (BPL) | 32 | 0 | 32 | 0 |
|  | A/bovine/OH/B24OSU-432/2024 (BPL) | 32 | 0 | 32 | 0 |
| 3* | A/CA/04/2009 (Live) | 16 | 0 | 0 | 12 |
|  | A/bovine/OH/B24OSU-432/2024 (Live) | 24 | 0 | 24 | 3 |

In conclusion, four different assay formats indicate that A/bovine/OH/B24OSU-432/2024 (H5N1) exhibits “*avian*” type virus receptor specificity. This agrees with the observation of inefficient transmission of the virus between ferrets^7^ and also between humans. Furthermore, A/bovine/OH/B24OSU-432/2024 (H5N1) maintains HA Q226 and G228 which has been associated with 2,3-sialoside receptor specificity. It is unclear as to why our data differ from those previous studies of Eisfeld *et al*. who did detect similar binding of a bovine H5N1 virus to α2,3- and α2,6-linked sialosides. In the previous study, a solid phase binding assay was used in the absence of an NA inhibitor, and it is possible that the greater activity of NA for α2,3-sialosides diminishes these receptors thereby skewing the binding data to residual α2,6-sialoside receptors. Of note, some binding to α2,6-sialoside receptors has been reported for naturally occurring avian H5 viruses^8^. As discussed, the ferret model has suggested that a complete switch from α2,3 to α2,6-sialoside binding is essential to confer efficient transmission^16,17^. It is likely that the same is necessary for A/bovine/OH/B24OSU-432/2024 adaptation to humans. The mammary tissue of cattle display mainly α2,3-sialosides^18^ and opportunities for adaptation to human-type receptor at this anatomical site are expected to be low. Deep mutational scanning of pseudoviruses has indicated that clade 2.3.4.4b HA can in principle acquire human virus receptor binding specificity^19^, and, thus, while we find no current evidence, ongoing effort must be placed on continued monitoring for this trait.

## Author contributions

P.C., R.J.W., R.P.V., S.M.T., and G.J.B. conceived the study. P.C., C.K.P., S.D.R., J.D.S., R.P.V., A.K., and T.J. performed the experiments. P.C., C.K.P., S.D.R., J.D.S., R.J.W., R.P.V., S.M.T., and G.J.B. analyzed the data and interpreted the results. A.S.B. isolated A/bovine/OH/B24OSU-432/2024 and A.H.E. developed reagents. P.C., R.P.V., S.M.T., and G.J.B. wrote the manuscript. R.J.W., S.M.T., and G.J.B. supervised research. All authors have given approval to the final version of the manuscript.

## Data and materials availability

All data is available in the main text or the supplementary materials. Sharing materials described in this work will be subject to standard material transfer agreements.

## Code availability

The script for Microsoft Excel Macro for batch processing glycan microarray data is uploaded to https://github.com/enthalpyliu/carbohydrate-microarray-processing.

