## Supplementary Information for "Receptor Binding Specificity of a Bovine A(H5N1) Influenza Virus"

**Table of contents** page

Figure S1. Characterization of *N*-glycan array using plant lectins S2

Figure S2. Solid phase assays for receptor binding specificities S3

Table S1. HA receptor binding site amino acid alignment of the H5 hemagglutinins S3

1. Virus production S4

2. Microarray printing and binding analysis S5

3. Biolayer interferometry binding assays S6

4. Solid-phase binding assay S7

5. Erythrocyte remodeling, virus titration and hemagglutination assay S7

6. References S9

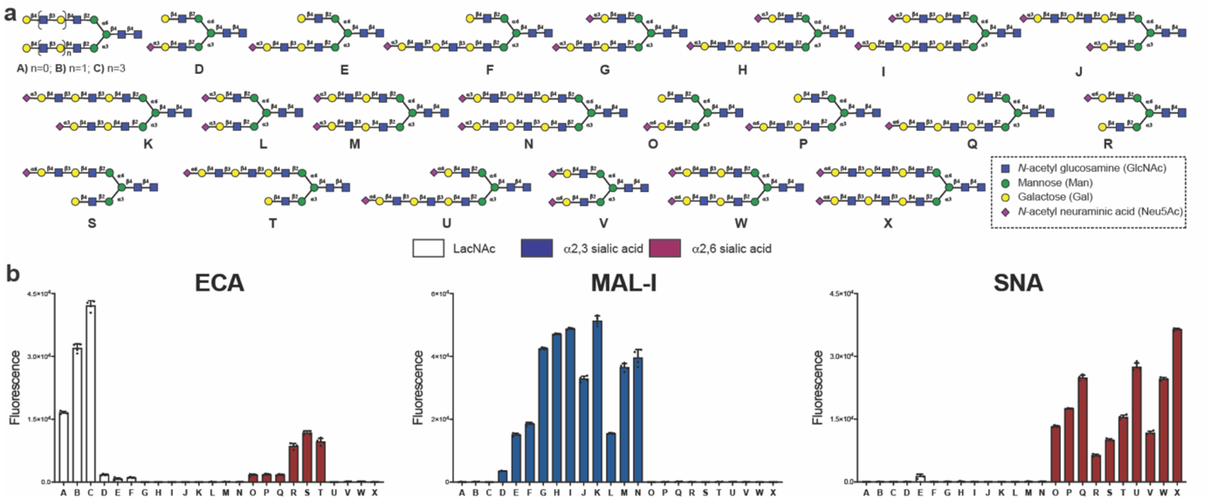


**Figure S1. Characterization of *N*-glycan array using plant lectins.** (a) Structures of *N*-glycans (**A**-**X**) printed on the microarray. (b) Glycan array binding analysis of the biotinylated plant lectins, *Erythrina cristagalli* agglutinin (ECA, specific for terminal Galactose), *Maackia amurensis* lectin I (MAL-I, specific for α2,3-linked Neu5Ac), and *Sambuca nigra* agglutinin (SNA, specific for α2,6-linked Neu5Ac). The lectins were detected using streptavidin-AlexaFluor 635 conjugate and fluorescence signals for each glycan are presented as mean ± SD (n = 4).

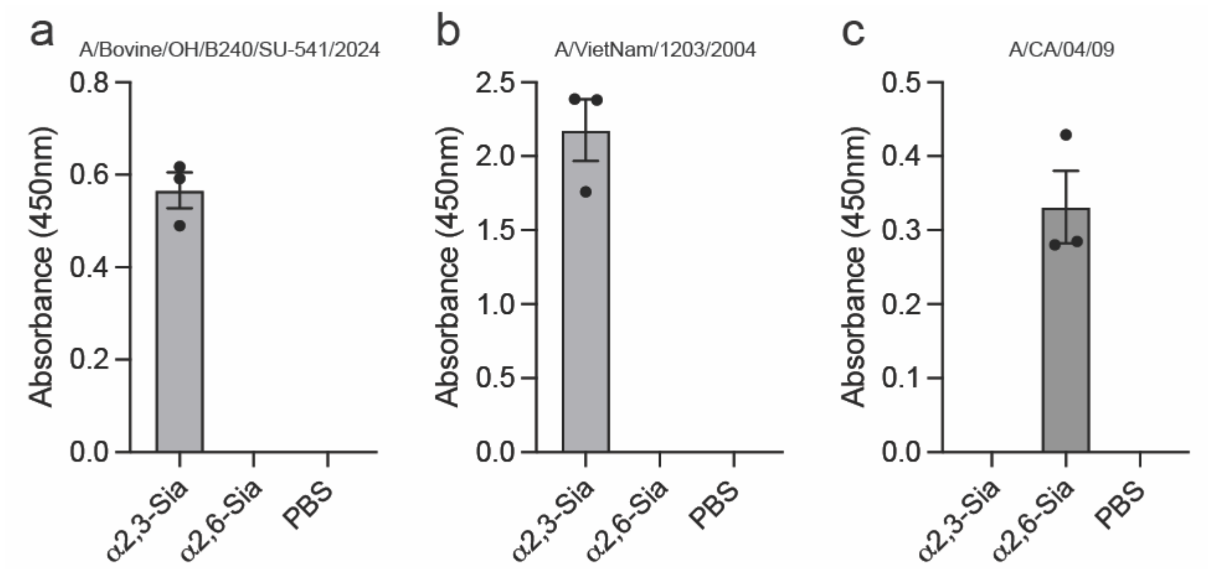


**Figure S2. Solid phase assays for receptor binding specificities.** The influenza A viruses (IAV) A/bovine/OH/B240SU-541/2024, A/Vietnam/1203/2004 and A/California/04/2009 (32 HA units) were captured on a fetuin coated plate and incubated with 3’-SLN-PAA-biotin (10 µg/mL), 6’-SLN-PAA-biotin (10 µg/mL) or PBS. Virus-bound biotinylated polymer was detected using horseradish peroxidase (HRP)-conjugated streptavidin. The absorbance ($A_{450}$) values for each treatment of all viruses were plotted using Prism 10 software (GraphPad Software, Inc.). The assay was performed in triplicate at least two times for each virus, bars (scatter dot plot) represent the mean ± SEM for each treatment.

| | 94 | 95 | 96 | 122 | 123 | 124 | 125 | 125a | 125b | 126 | 127 | 128 | 129 | 130 | 131 | 132 | 133 | 133a | 134 | 135 | 136 | 137 | 138 | 139 | 140 | 141 | 142 | 143 | 144 | 145 | 146 | 147 |
|---|---|---|---|---|---|---|---|---|---|---|---|---|---|---|---|---|---|---|---|---|---|---|---|---|---|---|---|---|---|---|---|---|
| A/VietNam/1203/2004 | C | Y | P | Q | I | I | P | K | S | S | W | S | S | H | E | A | S | L | G | V | S | S | A | C | P | Y | Q | G | K | S | S | F |
| A/dairy cattle/New Mexico/A240920343-93/2024 | . | . | . | . | . | . | . | . | . | . | . | P | N | . | . | T | . | . | . | . | . | A | . | . | . | . | . | . | A | P | . | . |
| A/bovine/OH/B24OSU-432/2024 | . | . | . | . | . | . | . | . | . | . | . | P | N | . | . | T | . | . | . | . | . | A | . | . | . | . | . | . | A | P | . | . |

| | 148 | 149 | 150 | 151 | 152 | 153 | 154 | 155 | 156 | 157 | 158 | 159 | 160 | 161 | 162 | 183 | 184 | 185 | 186 | 187 | 188 | 189 | 190 | 191 | 192 | 193 | 194 | 195 | 196 | 197 | 198 | 199 |
|---|---|---|---|---|---|---|---|---|---|---|---|---|---|---|---|---|---|---|---|---|---|---|---|---|---|---|---|---|---|---|---|---|
| A/VietNam/1203/2004 | F | R | N | V | V | W | L | I | K | K | N | S | T | Y | P | H | H | P | N | D | A | A | E | Q | T | K | L | Y | Q | N | P | T |
| A/dairy cattle/New Mexico/A240920343-93/2024 | . | . | . | . | . | . | . | . | . | . | . | D | A | . | . | . | . | S | . | N | . | E | . | . | . | N | . | . | K | . | . | I |
| A/bovine/OH/B24OSU-432/2024 | . | . | . | . | . | . | . | . | . | . | . | D | A | . | . | . | . | S | . | N | . | E | . | . | . | N | . | . | K | . | . | I |

| | 200 | 201 | 202 | 203 | 204 | 205 | 206 | 207 | 208 | 209 | 210 | 211 | 212 | 213 | 214 | 215 | 216 | 217 | 218 | 219 | 220 | 221 | 222 | 223 | 224 | 225 | 226 | 227 | 228 | 229 | 230 |
|---|---|---|---|---|---|---|---|---|---|---|---|---|---|---|---|---|---|---|---|---|---|---|---|---|---|---|---|---|---|---|---|
| A/VietNam/1203/2004 | T | Y | I | S | V | G | T | S | T | L | N | Q | R | L | V | P | R | I | A | T | R | S | K | V | N | G | Q | S | G | R | M |
| A/dairy cattle/New Mexico/A240920343-93/2024 | . | . | . | . | . | . | . | . | . | . | . | . | . | . | A | . | K | . | . | . | . | . | Q | . | . | . | . | R | . | . | . |
| A/bovine/OH/B24OSU-432/2024 | . | . | . | . | . | . | . | . | . | . | . | . | . | . | A | . | K | . | . | . | . | . | Q | . | . | . | . | R | . | . | . |

**Table S1. HA receptor binding site (RBS) amino acid alignment of the H5 hemagglutinins.** Alignment of the RBS residues with amino acid positions (94-230), including an extended 130-, 150-, and 220- loop and the 190-helix shown above the alignment, non-conserved residues highlighted in black, and the dots indicate conserved amino acids.

**1. Virus production**

*Virus production in cells*: Madin Darby Canine Kidney (MDCK) cells were maintained in Dulbecco's Modified Eagle Medium (DMEM; Corning Ref #10-013-CV) supplemented with 10% fetal bovine serum (FBS; Biowest Cat # S1400) and 1X antibiotics-antimycotic (A/A;GenDEPOT, Cat # 002-010) at 37°C with 5% $CO_2$ until reaching ~80% confluency. For propagation of A/California/04/2009, confluent MDCK cells were washed twice with 10 mL of sterile PBS and then a mixture of 10 mL DMEM, A/A, 1 mg/mL TPCK trypsin (Worthington, Cat # LS003740) and a 0.1 MOI of virus was added to the cells and incubated for 72 h at 37 °C with 5% $CO_2$. At 72 h post-infection, when cytopathic effect (CPE) was observed, the supernatant was collected. Cellular debris was removed by centrifugation (3000 g, 10 min, 4 °C) and the clarified supernatant containing the virus was aliquoted, and stored at -80°C.

*Virus production in eggs*: Ten-day old embryonated hens' eggs were inoculated with A/bovine/OH/B24OSU-432/2024, A/bovine/OH/B24OSU-541/2024, and A/Vietnam/1203/2004 at multiple dilutions and were incubated at 35 °C for 36 to 40 h. The eggs were candled for embryo viability beginning around 36 h and any non-viable embryonated eggs were chilled at 4 °C overnight. Allantoic fluid was dipped for HA and all positives were harvested. HA titer and sterility of the virus isolates were noted, and the stocks were stored at -80 °C. All virus isolation work and BPL inactivation was performed in approved Biosafety Level 3 enhanced (BSL3-enhanced) laboratories at St. Jude Children's Research Hospital (SJCRH) in Memphis, TN.

*BPL inactivation of viruses*: Each 1 mL of virus stock was inactivated by adding 1 μL of β-propiolactone (BPL) (Sigma, Cat# P5648, 10 mL). The tubes were inverted 10 times to thoroughly mix the virus and BPL and then poured into a new sterile tube. The BPL-inactivated viruses were stored at 4 °C for at least 72 h. Each BPL-inactivated virus was serially passaged in ten-day old embryonated hens' eggs as described for virus propagation. Virus inactivation was successful, and BPL inactivated samples were removed from BSL3-enhanced laboratories at SJCRH in Memphis, TN.

### 2. Microarray printing and binding analysis

*N*-glycans (**A**-**X**) with an α-amine at the reducing end asparagine moiety and *O*-glycans (**1**-**5**) with a *N*-terminal α-amine were printed on amine reactive, NHS-ester activated glass slides (NEXTERION® Slide H, Schott Inc.) using a Scienion sciFLEXARRAYER S3 non-contact microarray equipped with a Scienion PDC80 nozzle (Scienion Inc.). Individual samples were dissolved in sodium phosphate buffer (50 µL, 0.225 M, pH 8.5) at a concentration of 100 µM and were printed in replicates of 6 with a spot volume of ~ 400 pL at 20 °C and 50% humidity. Each slide has 24 subarrays in a 3x8 layout. After printing, slides were incubated in a humidity chamber for 8 h and then blocked for 30 min in a Tris buffer (pH 9.0, 50 mM) containing 5 mM ethanolamine at 50 °C. Blocked slides were rinsed with DI water, spun dry, and kept in a desiccator at room temperature for future use. For quality control, the printed slides were incubated with plant lectins *Erythrina cristagalli* agglutinin (ECA, specific for terminal Gal, Cat # B-1145, Vector Labs), *Maackia amurensis* lectin I (MAL-I, specific for α2,3-linked Neu5Ac, Cat # B-1315, Vector Labs), and *Sambuca nigra* agglutinin (SNA, specific for α2,6-linked Neu5Ac, Cat # B-1305, Vector Labs).

Binding was performed by incubating the slides with virus isolates (at 1:5 dilution) in TSM binding buffer (TSM-BB, Tris-HCl 20 mM pH 7.4, NaCl 150 mM, $CaCl_2$ 2mM, $MgCl_2$ 2mM, containing 1% BSA and 0.05% Tween-20) in the presence of a neuraminidase inhibitor (Oseltamivir carboxylate, OC, 10 µM) for 2 h at room temperature. Then, slides were washed by sequential dipping in TSM wash buffer (2 min, containing 0.05 % Tween 20), TSM buffer (2 min) and, water (2 x 2 min), followed by centrifugation. Next, slides were incubated with a premixed solution of 1A06 pan-HA stem antibody (5 µg/mL, from Dr. Ali Ellebedy, Washington University, MO)[1] and AlexaFluor 647-labeled goat anti-human IgG antibody (5 µg/mL, Jackson ImmunoResearch Cat #109-605-008) in TSMBB in the presence of OC (10 µM) for 1 h, followed by washing and drying. The biotinylated lectins were detected with streptavidin-AlexaFluor 635 conjugate (5 µg/mL, Thermo, Cat # S32364) (Extended data Fig. 1).

The slides were scanned using a GenePix 4000B microarray scanner (Molecular Devices) at the appropriate excitation wavelength with a resolution of 5 µM. Optimum gains and PMT values were employed for the scanning ensuring that all signals were within the linear range of the scanner's detector and there was no saturation of signals. The images were analyzed using GenePix Pro 7 software (version 7.2.29.2, Molecular Devices). The data were analyzed with a home written Excel macro. The highest and the lowest value of the total fluorescence intensity of the replicate spots were removed, and the remaining values were used to provide the mean value and standard deviation. The fluorescence values were plotted using Prism 10 software (GraphPad Software, Inc.), bars represent the mean ± SD for each treatment.

**3. Biolayer interferometry (BLI) binding assays**

A glycan functionalized biosensor was prepared by capturing a biotinylated glyco-polymer on a streptavidin coated biosensor (Octet® SAX Biosensors, part # 18-5117, Sartorius). Briefly, on a bio-layer interferometer (BLI) Octet Red 384 system (ForteBio, Pall life sciences), pre-hydrated (10 min) SAX biosensors in the assay buffer (Tris-HCl 20 mM pH 7.4, NaCl 150 mM, $CaCl_2$ 2 mM, $MgCl_2$ 2 mM, containing 1% BSA and 0.05% Tween-20) were dipped into wells of black-walled 384-well plate containing 100 μL of Lactose-PAA-biotin (5 μg/mL, Cat # OL139178, Biosynth), 3'-α-Sialyl-*N*-acetyllactosamine-PAA-biotin (5 μg/mL, 3'-SLN, Cat # OS137338, Biosynth) or 6'- α-Sialyl-N-acetyllactosamine-PAA-biotin (5 μg/mL, 6'-SLN, Cat # OS137337, Biosynth). Once saturation of the signal was observed, typically in 600 sec with a thickness reaching at 1 nm, the sensors were cleaned and stabilized by submerging in well containing assay buffer for 300 sec. Steps for loading sequence are as follows: baseline 60 sec, loading 600 sec, and washing 300 sec (shaking at 1000 rpm, temperature at 25 °C, for all steps).

For direct binding experiment, the virus isolates A/bovine/OH/B24OSU-432/2024 (at 1:5 dilution), A/Vietnam/1203/2004 (at 1:2 dilution) and A/California/04/2009 (at 1:2 dilution) were suspended at indicated dilution in the assay buffer (100 μL, final volume) containing Oseltamivir carboxylate (OC, 10 μM). The glycan functionalized sensors were first dipped in assay buffer to obtain a baseline, then in a virus suspension for association step, followed by dissociation in assay buffer lacking OC. Steps for binding assay are as follows: baseline 60 sec, association 1800 sec (3000 sec for A/bovine) and dissociation 1800 sec (3000 sec for A/bovine) (shaking at 1000 rpm, temperature at 25 °C, for all steps).

Data acquisition and data evaluation was performed using ForteBio Data Acquisition 11.1 software and ForteBio Data Analysis 11.1 software, respectively. The raw data for response curves were exported as an excel sheet (Microsoft Office Excel) and curves were flipped i.e., corrected to give positive response (by multiplying responses with -1). Curves depicting virus binding (thickness, nm) *vs.* time (sec) were plotted using Prism 10 software (GraphPad Software, Inc.). The assay was performed at least two times for each virus.

**4. Solid-phase binding assay**

To determine the receptor specificity of viruses, solid-phase binding assays were performed following the literature report[2]. Briefly, influenza viruses, A/bovine/OH/B24OSU-541/2024, A/Vietnam/1203/2004 and A/California/04/2009 (32 HA units in TBS, 50 µL) were added to a clear flat-bottom 96-well microtiter plate (NUNC MaxiSorp, Thermo, Cat # 439454) which was pre-coated with fetuin (from fetal bovine serum, Sigma Aldrich, Cat # F2379) and incubated at 4 °C overnight. Next day, the wells were aspirated, washed (PBS, 200 µL, 2-times), and blocked (0.1% BSA in PBS) for 2 h at room temperature. The wells were aspirated, washed with washing solution (cold PBS containing 0.01% Tween-20, 200 µL, 3 times), and 10 µg/mL solutions of Lactose-PAA-biotin, 3'-SLN-PAA-biotin, 6'-SLN-PAA-biotin in reaction buffer (PBS containing 0.1% BSA, 0.02% Tween-20 and 2 µM of OC) (50 µL each) were added and plate was incubated at 4 °C. After 2 h, the plate was washed with washing solution and then incubated with horseradish peroxidase conjugated streptavidin in reaction buffer (50 µL, 1:500 dilution, BioLegend 405210) for 1 h at 4 °C. After washing, the wells were incubated with TMB one component substrate solution (100 µL, SouthernBiotech, Cat # 0410-01) for 0.5 h at room temperature, then reaction was stopped by addition of 1M $H_2SO_4$ (50 µL) and the absorbance ($\lambda$ = 450 nm, POLARstar Optima, BMG Labtech) was measured. The absorbance was corrected for blank (no virus and no polymer) well and lactose glycopolymer (non-specific interaction) well in Microsoft office Excel and plotted using Prism 10 software (GraphPad Software, Inc.) (Extended data Fig. 2). The assay was performed in triplicate at least two times for each virus.

**5. Erythrocyte remodeling, virus titration and hemagglutination assay**

Turkey erythrocytes were obtained from the Poultry Diagnostic Research Center (PDRC) at the University of Georgia (UGA), Athens, USA, following protocol reviewed and approved by the Institutional Animal Care and Use Committee (IACUC) at the UGA. The erythrocytes were washed twice with sterile PBS (50 mL, pH 7.4, Corning, Cat # 21-040-CV) by centrifugation (300 g, 10 min) and diluted to a working concentration of 1% in PBS (v/v). For glycan remodeling, four separate preparation of 1.25 mL of 1% (v/v) erythrocytes in PBS were pelleted by centrifugation (300 g, 5 min) and resuspended in 62.5 µL of PBS. One preparation was resuspended in 1.25 mL of PBS (0.5% v/v) to serve as the untreated control and stored at 4 °C until further use. Three preparations of erythrocytes were suspended further in a 5:1 ratio of

1U/mL of *Arthrobacter ureafaciens* neuraminidase (New England Biolabs, Cat # P0722L) in a reaction buffer (PBS containing 0.1M $CaCl_2$) in a final reaction volume of 122.5 µL and incubated for 1 h at 37 °C on an orbital shaker. Following incubation, the cells were washed twice with PBS (1 mL) by centrifugation, and resuspended in PBS (50 µL, containing 1% BSA, Fisher Bioreagents, Cat # BP9704-100). Resialylation of erythrocytes in a final volume of 75µL with α2,3-linked Neu5Ac or α2,6-linked Neu5Ac was performed with α2,3-sialyltransferase (PMST1 M144D, 1 µg)[3] or α2,6-salyltransferase (ST6Gal1, 1 µg)[4], respectively in the presence of CMP-Neu5Ac (3.75 µL, 30 mM stock in PBS containing 1% BSA) and incubated for 4 h at 37 °C on an orbital shaker. The erythrocytes were again washed twice with PBS (1 mL) and diluted to a working concentration of 0.5% (v/v) in PBS (containing 1% BSA) for hemagglutination assays.

*Virus titration*: To a clear V-bottom 96-well microtiter plate (Greiner Bio-One, Cat # 651101), PBS (50 µL) was added to each well in a row. Next, 50 µL of either A/California/04/2009, A/Vietnam/1203/2004 (BPL inactivated), or A/bovine/OH/B24OSU-432/2024 (BPL inactivated) were added to the first well in the row and serially diluted two-fold across the plate. A suspension of turkey erythrocytes (0.5% v/v) was prepared in PBS and 50 µL were added to each well. The plate was incubated at room temperature, after 0.5 h, the plate was tilted at 90° to observe hemagglutination. The reciprocal of the highest dilution with hemagglutination was reported as the hemagglutination assay (HA) titer for the virus.

*Hemagglutination assay*: Viruses were diluted to an HA unit (HAU) of 32, and the HA was repeated to confirm titration. To perform the HA with remodeled erythrocytes, viruses with an HAU of 32 were diluted 2-fold in PBS across a V-bottom 96-well microtiter plate for a final volume of 50 µL. Next, 50 µL of a 0.5% suspension of untreated, neuraminidase treated, α2,3-sialic acid remodeled, or α2,6-sialic acid remodeled erythrocytes were added to each virus. After 0.5 h incubation at room temperature, the plate was tilted 90° to observe the absence or presence of hemagglutination at the dilution corresponding to a 32 HAU for each virus.
